# Endogenous retroviruses transcriptional modulation after severe infection, trauma and burn

**DOI:** 10.1101/433029

**Authors:** Olivier Tabone, Marine Mommert, Camille Jourdan, Elisabeth Cerrato, Matthieu Legrand, Alain Lepape, Bernard Allaouchiche, Thomas Rimmelé, Alexandre Pachot, Guillaume Monneret, Fabienne Venet, François Mallet, Julien Textoris

## Abstract

Although human endogenous retroviruses (HERVs) expression is a growing subject of interest, no study focused before on specific endogenous retroviruses loci activation in severely injured patients. Yet, HERV reactivation is observed in immunity compromised settings like some cancers and auto-immune diseases. Our objective was to assess the transcriptional modulation of HERVs in burn, trauma and septic shock patients. We analyzed HERV transcriptome with microarray data from whole blood samples of a burn cohort (n=30), a trauma cohort (n=105) and 2 septic shock cohorts (n=28, n=51), and healthy volunteers (HV, n=60). We described expression of the 337 probesets targeting HERV from U133 plus 2.0 microarray in each dataset and then we compared HERVs transcriptional modulation of patients compared to healthy volunteers. Although all 4 cohorts contained very severe patients, the majority of the 337 HERVs was not expressed (around 74% in mean). Each cohort had differentially expressed probesets in patients compared to HV (from 19 to 46). Strikingly, 5 HERVs were in common in all types of severely injured patients, with 4 being up-modulated in patients. We highlighted co-expressed profiles between HERV and nearby gene as well as autonomous HERV expression. We suggest an inflammatory-specific HERV transcriptional response, and importantly, we introduce that the HERVs close to immunity-related genes might have a role on its expression.

## Introduction

Human Endogenous Retroviruses (HERVs) are former exogenous retroviruses which have infected germinal cells and became integrated in our genome million years ago (Young, Stoye, and Kassiotis 2013). These rare events happened several times in evolution. As retrotransposons, they are able to duplicate across the genome and they represent today more than 8% of our genome. Each insertion therefore led to distinct groups or families, each including multiple copies. Current classification annotates around 100 such groups.

HERV loci initially shared a common structure with exogenous retroviruses: internal protein coding regions (*gag, pro, pol, env*) flanked by two identical Long Terminal Repeats (LTRs). The accumulation of mutations and recombination events during evolution made most of these elements incomplete and defective for replication. Indeed, most of HERVs in our genome are now solo LTRs (Young, Stoye, and Kassiotis 2013) resulting from recombination between 5’ and 3’ proviral LTRs. LTRs are critical elements that control viral gene expression either as promoters, enhancers or as polyadenylation signals. When inserted upstream, within or downstream of a “conventional” protein coding gene, LTRs can modulate its expression pattern (Cohen, Lock, and Mager 2009; Isbel and Whitelaw 2012). For example, the presence of intronic LTR can result in novel transcripts, by providing alternative promoters, enhancers or polyadenylation signals, or by altering RNA splicing (Jern and Coffin 2008; Mager et al. 1999; Dunn and Mager 2005). Very few is known about of the transcriptional modulation of such elements in pathological contexts but in cancers (like testicular cancer (J. Gimenez et al. 2010) or colorectal cancer (Pérot et al. 2015)) and auto-immune diseases (like multiple sclerosis (Laska et al. 2012; Balada, Vilardell-Tarrés, and Ordi-Ros 2010; Madeira et al. 2016)).

Few studies focused on HERVs reactivation in acute inflammatory contexts. In mice, modulation of HERVs expression has been shown to be quite specific, with signatures related to pathogen-associated molecular pattern (PAMPs) (Young et al. 2012). In human, LPS or PMA stimulations of myeloid cells revealed an increase expression of four HERVs families (Johnston et al. 2001). In vivo, HERVs expression has been detected in the plasma and whole blood samples of burn patients (Y.-J. Lee et al. 2013; K.-H. Lee et al. 2014) although the studies focused on whole HERVs families, not on specific loci. Studying HERV transcriptome modulation after severe inflammatory injuries could help to better understand pathological states of patients.

After severe injuries like septic shock, burn or trauma, leading to an important inflammatory response, we and others have shown that the blood transcriptome is highly modulated, with early and profound changes in adaptive and innate immune responses (Plassais et al. 2017; Xiao et al. 2011). Moreover in these contexts, viral reactivation is often observed, especially for Herpes Viruses (Ong et al. 2017; Textoris and Mallet 2017). This reactivation is associated with an immunosuppressive state (Walton et al. 2014). We therefore hypothesize that HERV, like latent viruses, may reactivate and be transcribed in vivo after inflammatory injuries. Given that several groups showed that some probes of commercial whole genome microarray do target HERV loci (Young, Mavrommatis, and Kassiotis 2014; Reichmann et al. 2012) (such as Affymetrix U133 plus 2), we retrospectively explored microarray datasets obtained in our lab to study the HERV transcriptome modulation in various contexts of injuries *in vivo*.

## Material and Methods

### Patients and sample collection

#### Microarray analyzed cohort

##### Burns cohort

30 severe burn patients admitted at Hospices Civils de Lyon, France (HCL) were included in a placebo-controlled, randomized, double-blind study assessing the efficacy of hydrocortisone administration on burn shock duration. Inclusion / exclusion criteria, clinical description and ethical considerations of the cohort have been previously published elsewhere (Venet et al. 2015; Plassais et al. 2017). Thirteen healthy volunteers were also recruited within Hospices Civils de Lyon to serve as controls for the transcriptional study. Whole blood samples were collected at inclusion (severe shock, before any treatment, Day 1) and in the following days (around day 2 (D2), day 5 (D5) and day 7 (D7) after inclusion).

##### Traumas cohort

105 patients with severe trauma were admitted at HCL. Briefly, patients were included when they were under mechanic ventilation, with an Injury Severity Score (ISS) over 25 and were at least 18 years old. Inclusion / exclusion criteria and ethical considerations of the cohort have been previously published elsewhere (Gouel-Chéron et al. 2015). The main clinical variables are summarized on Table S1. Samples were collected at day 1 (D1) or day 2 (D2) after trauma. Data from 22 healthy volunteers were also used to make comparisons with patients (identical with septic shock cohort 2).

##### Septic shock cohort 1 (SS1)

28 septic shock patients and 25 HV admitted into 2 ICUs of HCL were included in this study to explore the early transcriptome modulation after septic shock. Inclusion / exclusion criteria, clinical description and ethical considerations of the cohort have been previously published elsewhere (Cazalis et al. 2014). The first blood sample was collected at the onset of shock (i.e., within 30 min after the beginning of vasoactive treatment, D0) and at day 1 (D1) and day 2 (D2) after shock.

##### Septic shock cohort 2 (SS2)

51 septic shock patients admitted to two Intensive Care Units (ICU) of HCL and 22 HV were included in a prognostic biomarker study. Inclusion / exclusion criteria, clinical description and ethical considerations of the cohort have been previously published elsewhere (Venet et al. 2017). Samples were collected at day 1 (D1), day 2 (D2) and day 3 (D3) after shock.

#### RT-qPCR validation cohorts

##### Patients

Subset of cohorts used for microarray analysis were used for validation cohort: 10 burn samples at D1, 10 traumas samples at D1, 10 SS1 samples at D1, 10 SS2 samples at D1. Each subset was matched with its corresponding cohort on: Age, sex and Total Burn Surface Area (TBSA) for burns - Sex, Sepsis at D7 and Death at D28 for traumas - Age, sex and SAPS II for SS1 - Age, Sex and Death at D28 for SS2.

##### Healthy Volunteers

Whole blood samples were purchased from the Etablissement Français du Sang (n=12). The mean age of HV is 56, with a standard error of 9. According to the standardized procedure for blood donation, written informed consent was obtained from healthy volunteers (HVs) and personal data for blood donors were anonymized at time of blood donation and before blood transfer to a research lab.

#### Flow cytometry validation cohort

##### Burns

Whole blood samples (EDTA tubes) from 13 burn patients sampled at D1 and D7 and admitted in Edouard Herriot hospital at Lyon, France were recruited as part of the EARLYBURN study (NCT02940171). Patients were aged from 21 to 84 (mean = 53), 12 men. The mean TBSA was 33% (from 20% to 52%). All samples from these patients were used for CD300LF protein analysis, and 7 of these 13 patients were used for *CD55* protein analysis.

##### Septic shocks

Whole blood samples (EDTA tubes) from 22 septic shock patients sampled at D1/D2, D3/D4/D5 and D6/D7/D8 after shock and admitted in Edouard Herriot hospital at Lyon, France were recruited as part of IMMUNOSEPSIS study (NCT02803346). Patients were aged from 23 to 81 (mean = 68), 16 men. Eleven samples were used for CD300LF protein analysis and 11 other samples for *CD55* protein analysis.

##### Healthy volunteers

Whole blood samples (EDTA tubes) were purchased from the Etablissement Français du Sang (n=18). Donors were aged from 21 to 63 (mean = 50), 12 men and 6 women. They were age-matched with burn and septic shock cohorts. According to the standardized procedure for blood donation, written informed consent was obtained from healthy volunteers (HVs) and personal data for blood donors were anonymized at time of blood donation and before blood transfer to a research lab.

### RNA extraction and microarrays

Total RNA was extracted with PAXgene™ Blood RNA kit (PreAnalytix, Hilden, Germany). Whole blood from PAXGene™ tubes was preferred to either buffy coat or PBMCs to ensure reproducibility and avoid missing samples within the context of a clinical study. RNA integrity was assessed using Agilent 2100 Bioanalyser (Agilent Technologies, Waldbrom, Germany) and Lab-on-chip RNA 6000 Nano Assay (Agilent Technologies). Double-stranded cDNA was prepared from total RNA and an oligo-dT primer using GeneChip One-Cycle cDNA Synthesis Kit (Affymetrix, Santa Clara, United States). Three μg labeled cRNA were hybridized onto Human Genome U133 Plus 2.0 GeneChips (Affymetrix), revealed and washed using FS450 fluidic station. GeneChips were scanned using a 5G scanner (Affymetrix) and images (DAT files) were converted to CEL files using GCOS software (Affymetrix).

### Microarray analysis

Microarray data are available on the Gene Expression Omnibus (GEO) website for Burn [GEO:GSE77791], SS1 [GEO:GSE57065] and SS2 [GEO:GSE95233] cohorts. The preprocessing methods were comparable in all datasets. Microarray normalization and statistical analysis were performed using R/Bioconductor (R v3.2.3). Quality assessment was performed through simpleaffy (v2.46.0) (Wilson and Miller 2005). After removing outlier samples the raw data were normalized, adjusted for background noise and summarized using the GCRMA (Guanine Cytosine Robust Multi-Array) algorithm with default parameters (Wu and Irizarry 2005). COMBAT algorithm (Johnson, Li, and Rabinovic 2007) was used to remove batch effect on Burn and Trauma cohorts. The 337 probesets from the U133 Plus2.0 microarray targeting HERVs have been identified and selected as described elsewhere (Young, Mavrommatis, and Kassiotis 2014; Reichmann et al. 2012).

All the analysis were made with R (3.2.3). The differential expression analysis was performed with Limma package (3.26.9) (Ritchie et al. 2015). A probeset was considered significantly statistically differentially expressed between two conditions when absolute log2 Fold Change was higher than 0.5 and adjusted P-values (Benjamini-Hochberg correction (Benjamini and Hochberg 1995)) lower than 0.01.

### Reverse transcription and quantitative PCR

RNA from the cohorts, according to the above criteria, and new RNA from HV were selected. RNA concentration was determined using Quant-iT RNA, BR assay on Qubit (Life Technologies, Chicago, Ilinois, United States). RNA integrity was assessed with the RNA 6000 Nano Kit on a Bioanalyzer (Agilent Technologies, Santa Clara, California, United States). Samples with RNA integrity number ≤ 6 were excluded due to poor quality RNA. Total RNA was reverse transcribed in complementary DNA (200ng in a final volume of 20 μL) using QuantiTect Reverse Transcription kit (Qiagen) as recommended by the manufacturer. The expression levels of genes (*CD55, CD300LF, SLC8A1, NFE4, PTTG1IP* and *HPRT1* as reference gene) and associated HERVs were quantified using quantitative-real time polymerase chain reaction (qPCR). qPCR were performed on a LightCycler instrument using Light Cycler 480 Probes Master for the genes and reference genes and on SYBR Green I master for HERVs. Final volume of 20μL contains 0.5μM of primers. For genes, an initial denaturation step of 10min at 95°C followed by 45 cycles, 10 sec at 95°C, 29 sec annealing at 60°C, and 1 sec extension at 72°C, Taqman) was performed. For HERVs, an initial denaturation step of 5min at 95°C followed by 45 cycles of a PCR protocol (10 sec at 95°C, 15sec at 55°C and 15sec at 72°C, SYBR Green program), melting curve protocol was performed. The Second Derivative Maximum Method was used with the LightCycler software (Release 1.5.1) to automatically determine the crossing point for individual samples. Standard curves were generated by using serial dilutions of cDNA standards prepared from purified PCR amplicons obtained with the corresponding primers (Table S2). Relative standard curves describing the PCR efficiency of selected targets were created and used to perform efficiency-corrected quantification with the LightCycler Relative Quantification Software. Targets expression normalization was performed using a selected housekeeping gene (hypoxanthine phosphoribosyltransferase 1 [HPRT1, (Friggeri et al. 2016)]), and results were expressed as normalized concentration ratio.

### Flow cytometry

#### Sampling and staining

The following antibodies were used: anti CD14-BV510, anti CD3– BV421 and anti CD56–PECy7 from BD Biosciences; anti CD300lf-PE from BD Biosciences or anti *CD55*-APC from Biolegend; anti CD16-APC from BD Biosciences or anti CD16-FITC from Beckman Coulter (Miami, FL) and PE Mouse IgG1, κ Isotype Control from BD Biosciences or APC Mouse IgG1, κ Isotype Control from R&D System. Red blood cell lysis was performed using Versalyse lysing solution (Beckman Coulter). *CD300LF* and *CD55* expression were measured using Navios flow cytometer (Beckman-Coulter). Results were analyzed with Kaluza software (Beckman-Coulter) expressed as Medians of Fluorescence Intensity (MFI).

### Statistics

Wilcoxon signed rank tests were done for RT-qPCR and flow cytometry results, by comparison between HV and each cohort of patients, for each target.

### Ethics approval and consent to participate

EDTA blood tubes were obtained from EFS (Etablissement Français du Sang) and used immediately. In accordance with EFS standardized procedures for blood donation, written no-objection was obtained from healthy volunteers to use the blood for the research and personal data for blood donors were anonymized before blood transfer to our research lab.

Protocols of the discovery and validation cohorts were approved by local ethics committees. Non-opposition to inclusion in the protocols was systematically recorded from patients or next of kin.

## Results

We studied the *in vivo* modulation of the HERV transcriptome in three clinical relevant models of acute inflammatory injury: a burn, a trauma and 2 septic shock cohorts. We analyzed expression from each cohort independently comparing patients with healthy volunteers. All cohorts included severely injured patients (Table 1). The 30 burn patients had a median total burn surface area (TBSA) of 70% and high severity scores (median Baux: 110, median Abbreviated Burn Severity Index (ABSI): 11). The 105 trauma patients had a median Injury Severity Score (ISS) score of 34 and a median Simplified Acute Physiology Score II (SAPSII) of 44. The 28 septic shocks from SS1 cohort had a median SAPSII of 45 and a median Charlson score of 2. The 51 patients from SS2 cohort had a median SAPSII of 51.

**Table 1:**
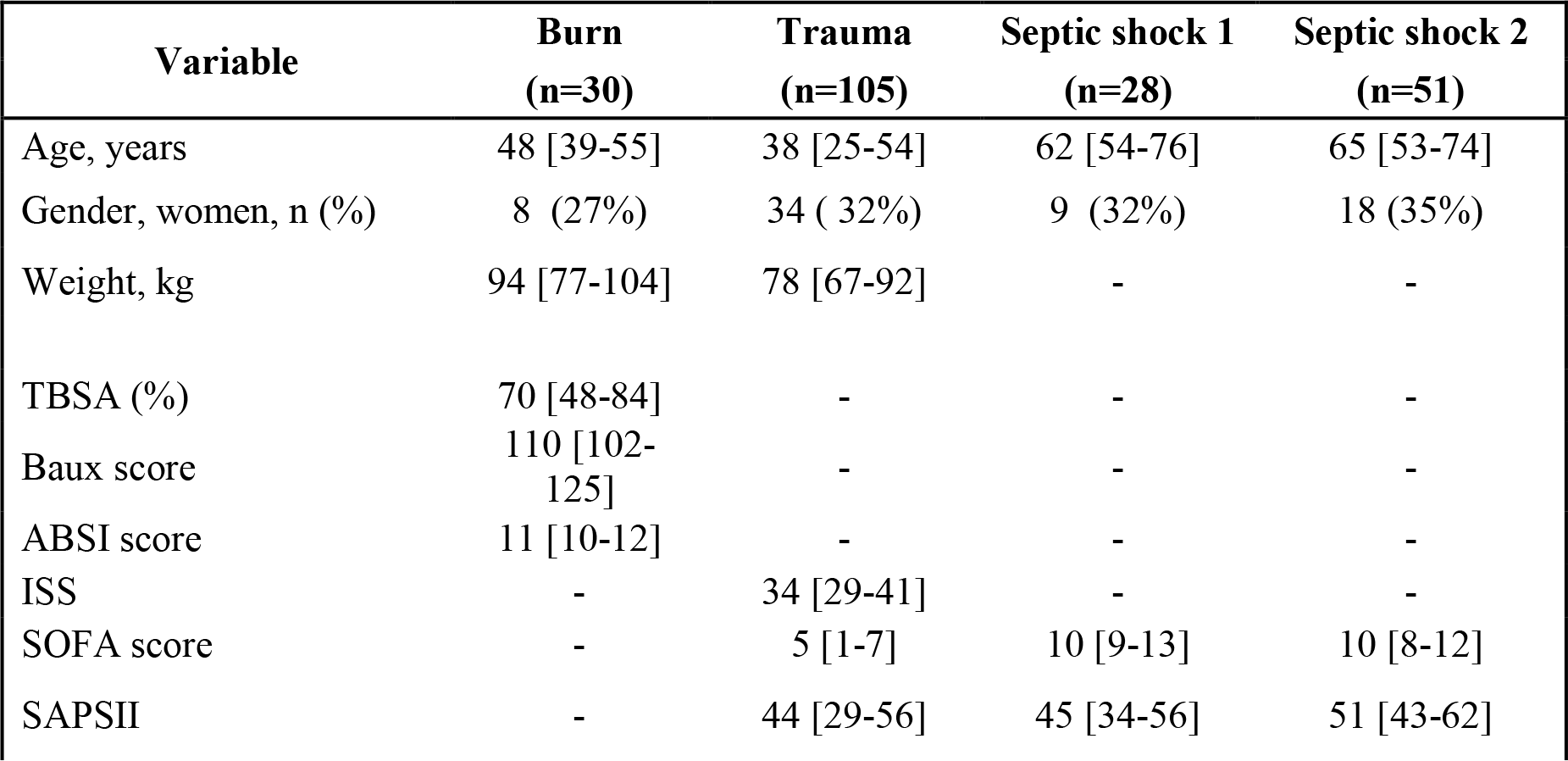
Patients characteristics of burn, trauma and septic shock cohorts included in microarray analyses. TBSA: Total Burn Surface Area; ISS: Injury Severity Score; ABSI: Abbreviated Burn Severity Index; SAPSII: Simplified Acute Physiology Score II.

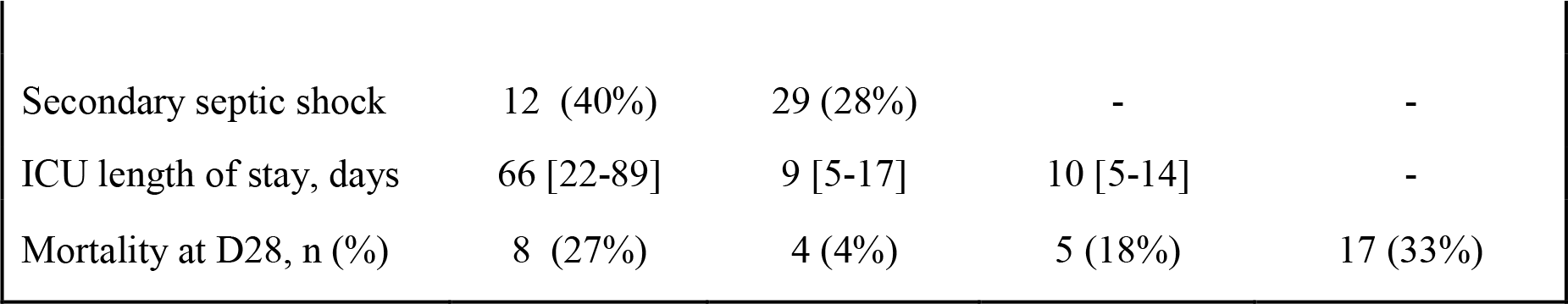

As previously published (Young, Mavrommatis, and Kassiotis 2014; Reichmann et al. 2012), we extracted data from 337 probesets targeting HERVs loci from the whole genome U133 plus 2.0 microarray datasets. Among them, a majority had low expression levels, within background levels (Figure 1). Based on hierarchical clustering analysis, 64 probesets (19%) were expressed (i.e. above background) for burns, 60 probesets (18%) for traumas, 164 for septic shock 1 (49%) and 63 for septic shocks 2 (19%). The 25% most variant probesets (n=84) across samples in each dataset revealed that several probesets were even highly expressed (Figure 2). In each dataset, the hierarchical clustering highlighted a clear difference between patients and HV, suggesting a modulation of HERV expression following injury. Interestingly, over these top 25% most variant probesets selected in each dataset (resulting of 127 distinct probesets), 44 (35%) were similarly modulated in the four datasets, and 102 (80%) in at least 2 datasets (Figure 3). In order to analyze the HERV transcriptome modulation associated with injury, we performed a supervised analysis comparing HERV expression in injured patients at D1 (admission) and HV, in each dataset separately. The comparison (accounting for multiple testing correction with absolute fold change higher than 1 and corrected p-value lower than 0.01) between burn patients and HV resulted in 19 differentially expressed HERVs (Figure 4A). The comparison between trauma patients and HV resulted in 27 differentially expressed HERVs (Figure 4B). The comparison between septic shock patients and HV resulted in 19 and 46 differentially expressed HERVs for cohorts 1 and 2 respectively (Figure 4C and D). Altogether, 56 different probesets targeting HERVs were differentially expressed among all 4 datasets, clearly discriminating HV from patients at ICU admission (Figure 5, Table S3). Taking into account the global profile for each probeset, 16 (28.6%) had higher expression in patients compared to HV and 40 (71.4%) were down-modulated in patients. Interestingly, 5 probesets were differentially expressed in all 4 datasets and 16 in at least 3 of them (Figure 6A). All 5 commonly modulated probesets had consistent expression profile across the 4 datasets. Four were over-expressed in patients compared to healthy volunteers (Figure 6B). The 5^th^ probeset, down-modulated in all datasets, maps at multiple locations in the genome and was not considered in further analyses. Among the 4 remaining modulated probesets, 1 HERV from ERV24B_Prim-int family (236982_at), is within 2kb from the *PTTG1IP* gene and 3 are within a gene. A HERV from LTR33 family (230354_at) is within an intron of *SLC8A1* gene. A HERV from MLT1H family (1556107_at) and one from LTR16B2 family (1559777_at) are located in the 3’UTR of *CD55* and *MIR3945HG* genes respectively (Table 2).

**Figure 1:**
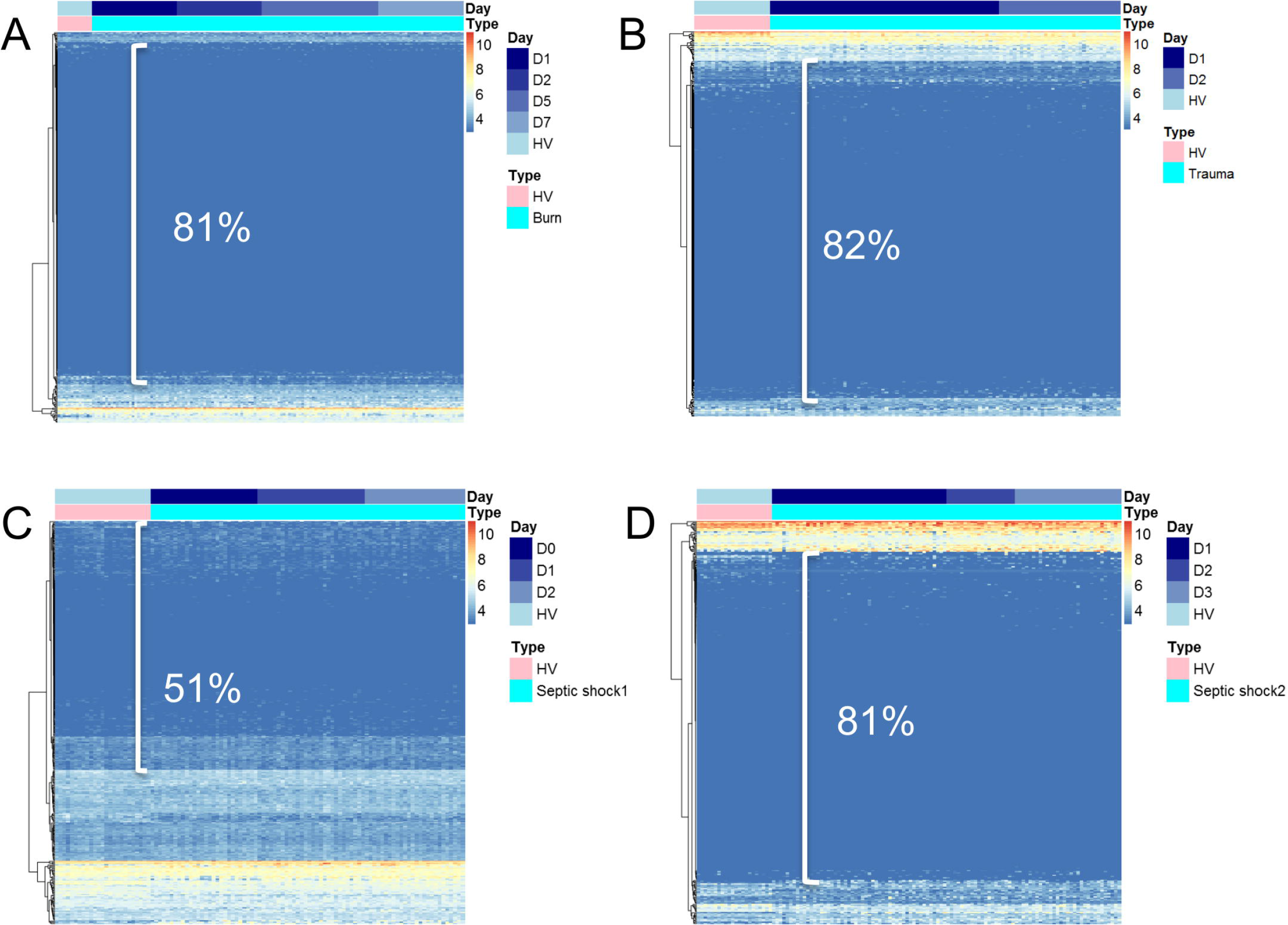
Heatmap representation of HERVs in three models of injury. Heatmap of the 337 probesets targeting HERVs in the four datasets: burn, trauma and 2 septic shock cohorts. Probesets are in rows and samples in columns. Samples are annotated (colored bars on the top) by type of samples (HV in pink, patients in cyan) and day after inclusion (blue scaled). Expression levels are color-coded from blue (low expression) to red (high expression). Similar patterns of expression are highlighted through hierarchical clustering of probesets (rows) with Euclidean distance and complete clustering method. **(A)** Expression levels in burns. **(B)** Expression levels in traumas. **(C)** Expression levels in septic shock 1. **(D)** Expression levels in septic shock 2. On each heatmap, the percentage of probesets with low intensity is shown.

**Figure 2:**
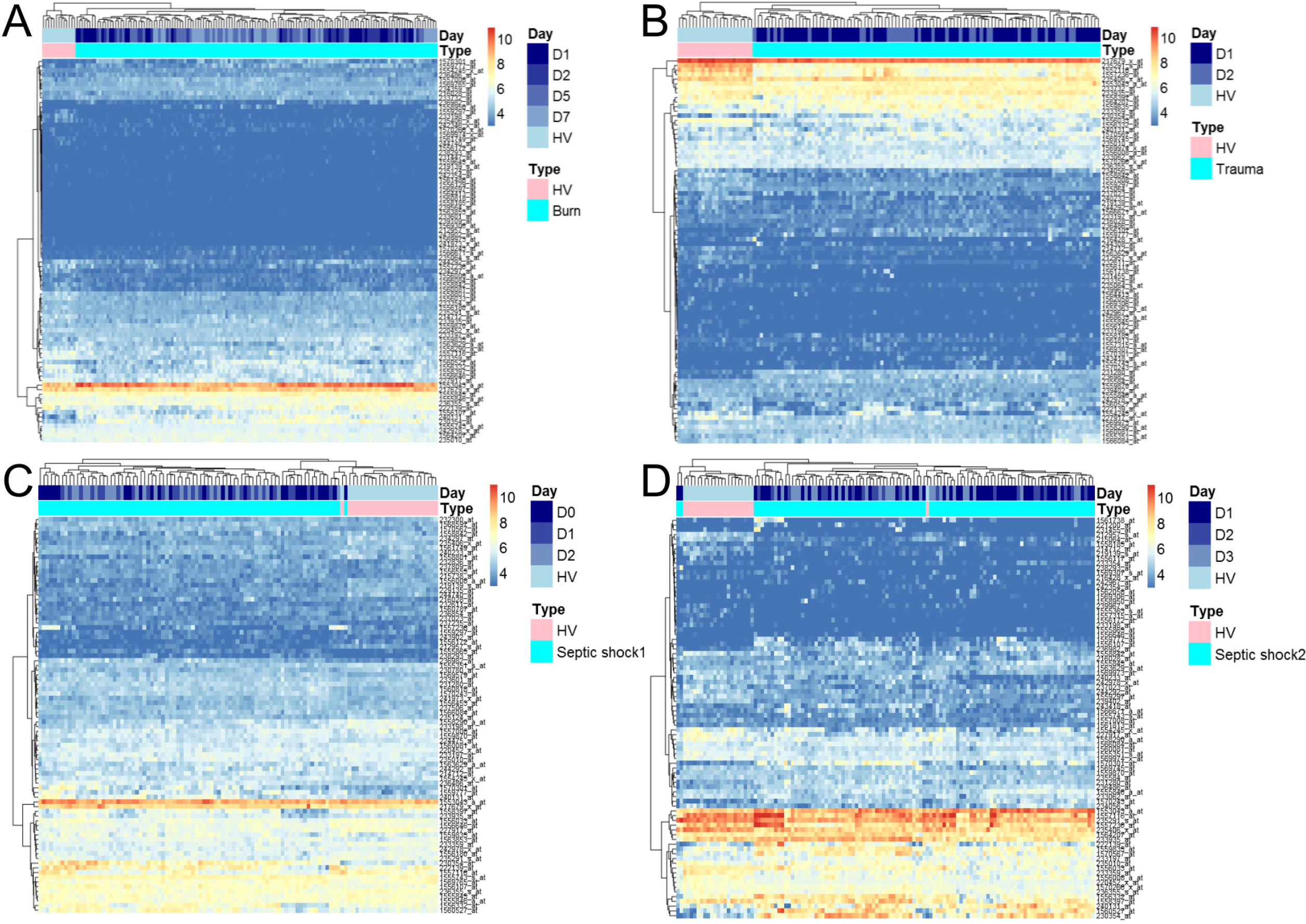
Heatmap representation of HERVs in three models of injury. Heatmap of the 25% most variant probesets targeting HERVs in the four datasets: burn, trauma and 2 septic shock cohorts. Probesets are in rows and samples in columns. Samples are annotated (colored bars on the top) by type of samples (HV in pink, patients in cyan) and day after inclusion (blue scaled). Expression levels are color-coded from blue (low expression) to red (high expression). Similar patterns of expression are highlighted through hierarchical clustering of probesets (rows) and samples (columns) with Euclidean distance and complete clustering method. **(A)** Expression levels in burn patients. **(B)** Expression levels in trauma patients. **(C)** Expression levels in septic shock 1 patients. **(D)** Expression levels in septic shock 2 patients.

**Figure 3:**
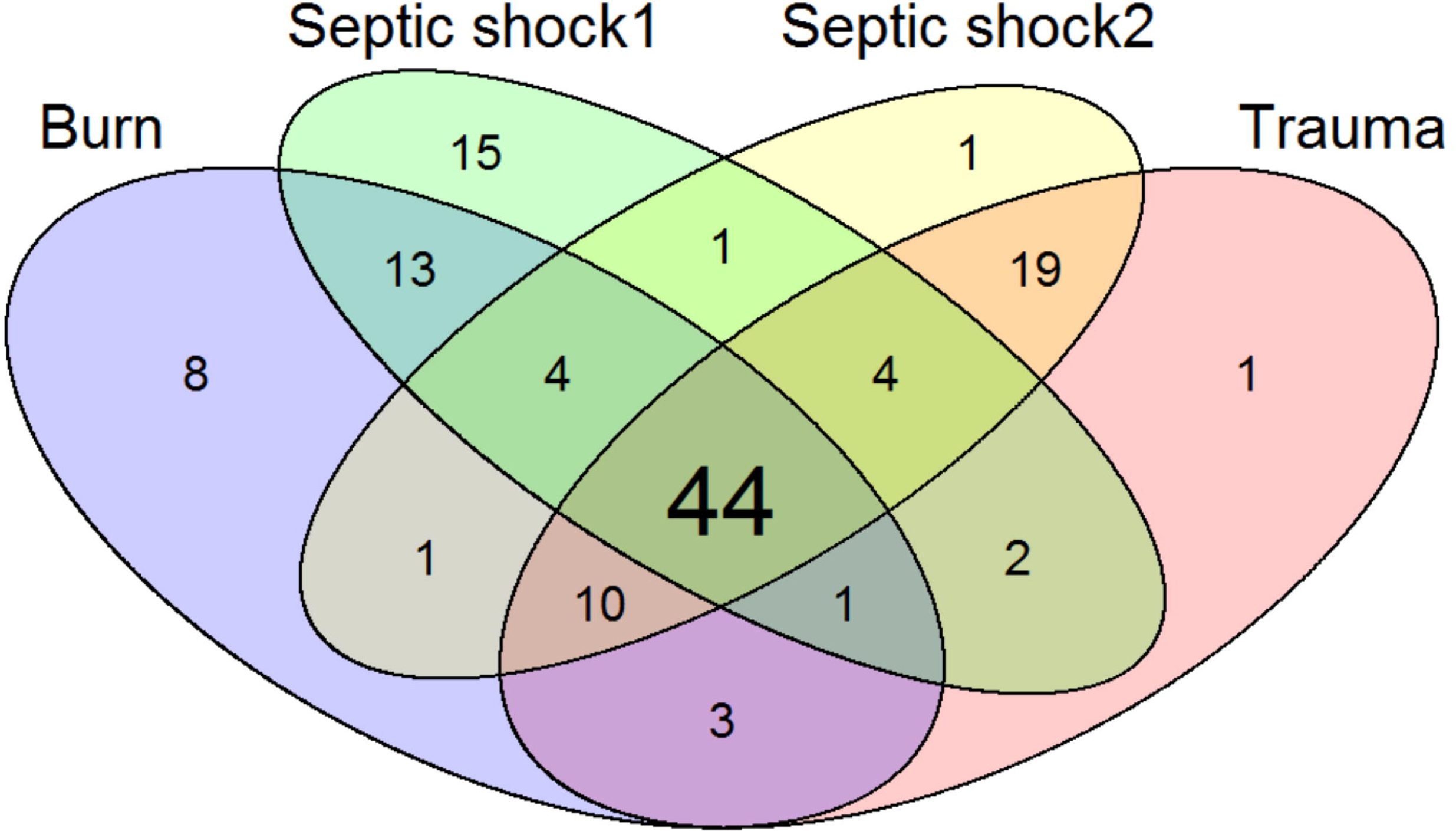
Most variant HERVs in severely injured patients. Venn diagram of the 84 most variant HERV probesets (25%) selected in each of the four datasets.

**Figure 4:**
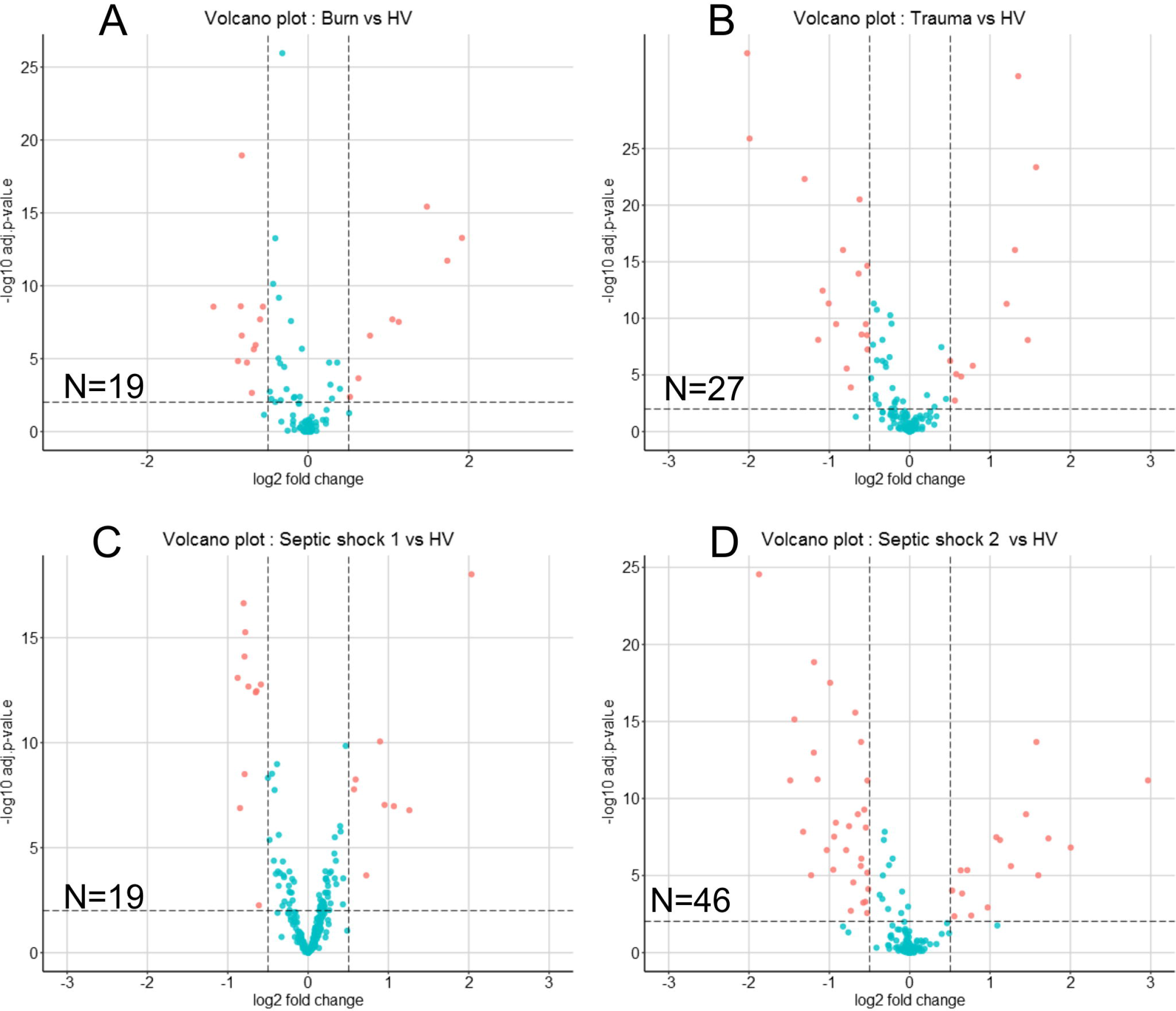
Volcano plots of differentially expressed HERVs. **(A)** in burn cohort. **(B)** in trauma cohort. **(C)** in septic shock cohort 1 and **(D)** in septic shock cohort 2. The x-axis represents the log2 fold change between patient and HV, the y-axis the −log10 of adjusted p-values. Each point represents a probeset targeting HERV, in red the statistically differentially expressed between patients at D1 and HV. On each volcano plot, the number indicates the number of differentially expressed probesets.

**Figure 5:**
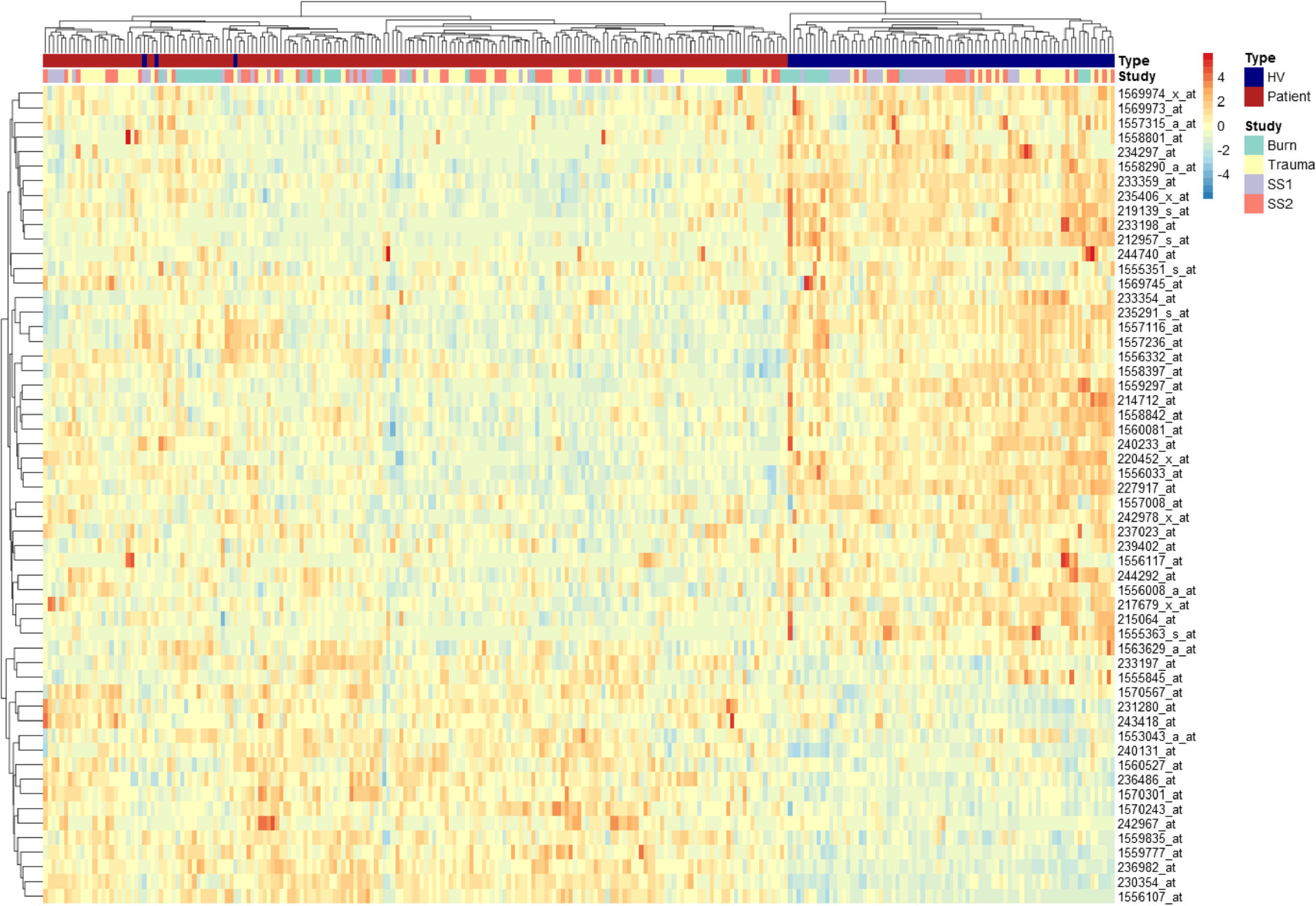
Heatmap representation of the modulated HERVs in severely injured patients at D1. Heatmap of the 56 differentially expressed probesets in at least 1 dataset. On the top bar, samples are color-coded in blue for HV and in red for Patients. On the bar below, samples are in green for Burn study, in yellow for Trauma study, in purple for Septic Shock 1 (SS1) study and in light red for Septic Shock 2 (SS2). Probesets are in rows and samples in columns. Expression levels from each cohort have been normalized (centered and reduced). Normalized expression levels are color-coded from blue (low expression) to red (high expression). Similar patterns of expression are highlighted through hierarchical clustering of probesets (rows) and samples (columns) with Euclidean distance and complete clustering method.

**Figure 6:**
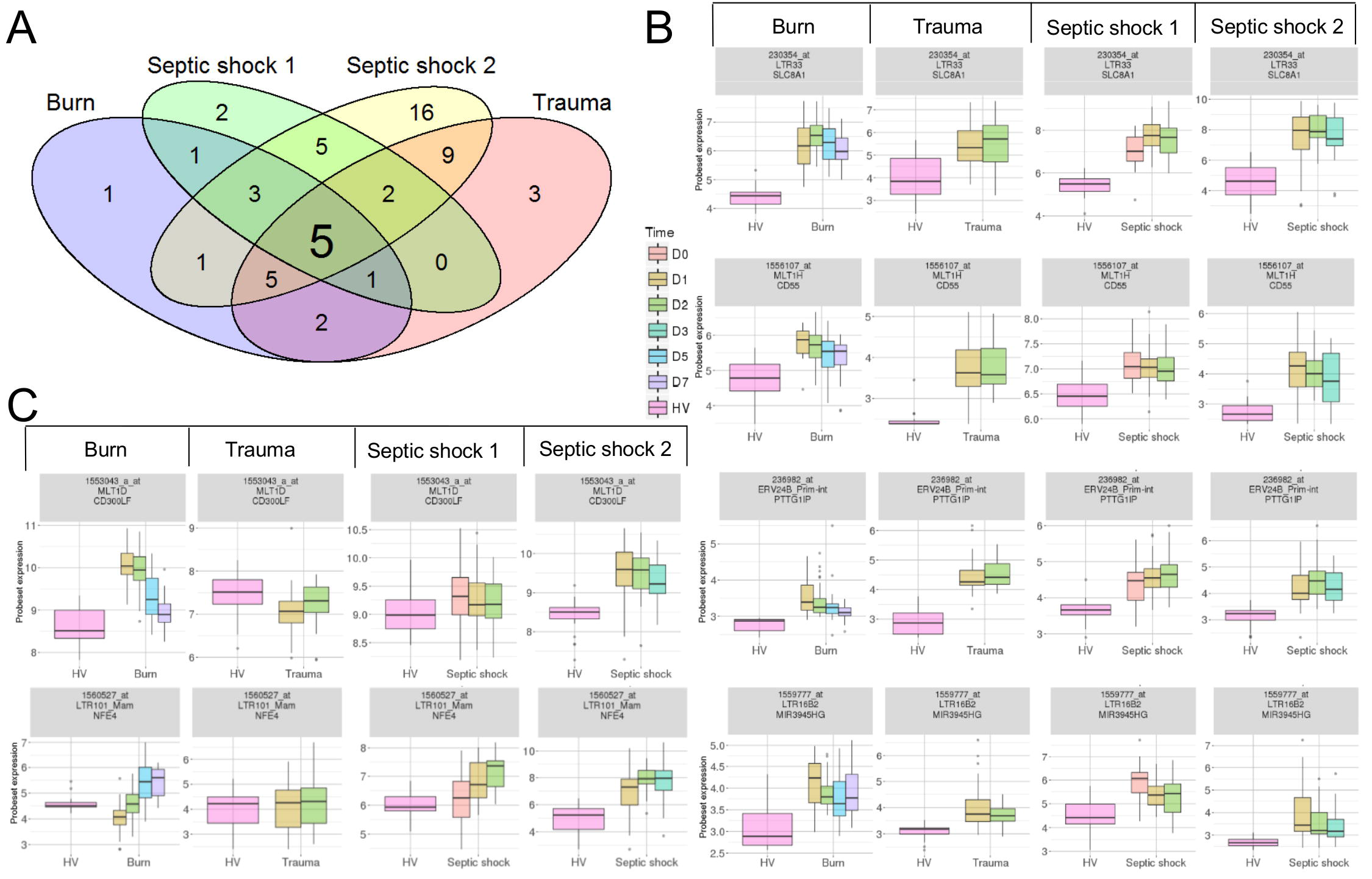
Differentially expressed HERVs in severely injured patients. **(A)** Venn diagram of differentially expressed HERVs for each dataset. **(B)** Expression profiles of commonly modulated probesets targeting HERVs in the 4 datasets. Boxes are color-coded by day after inclusion. **(C)** Expression profiles of 2 selected probesets targeting HERVs. For each graphic from top to bottom, title contains: probeset name, HERV name and closest gene.

**Table2:**
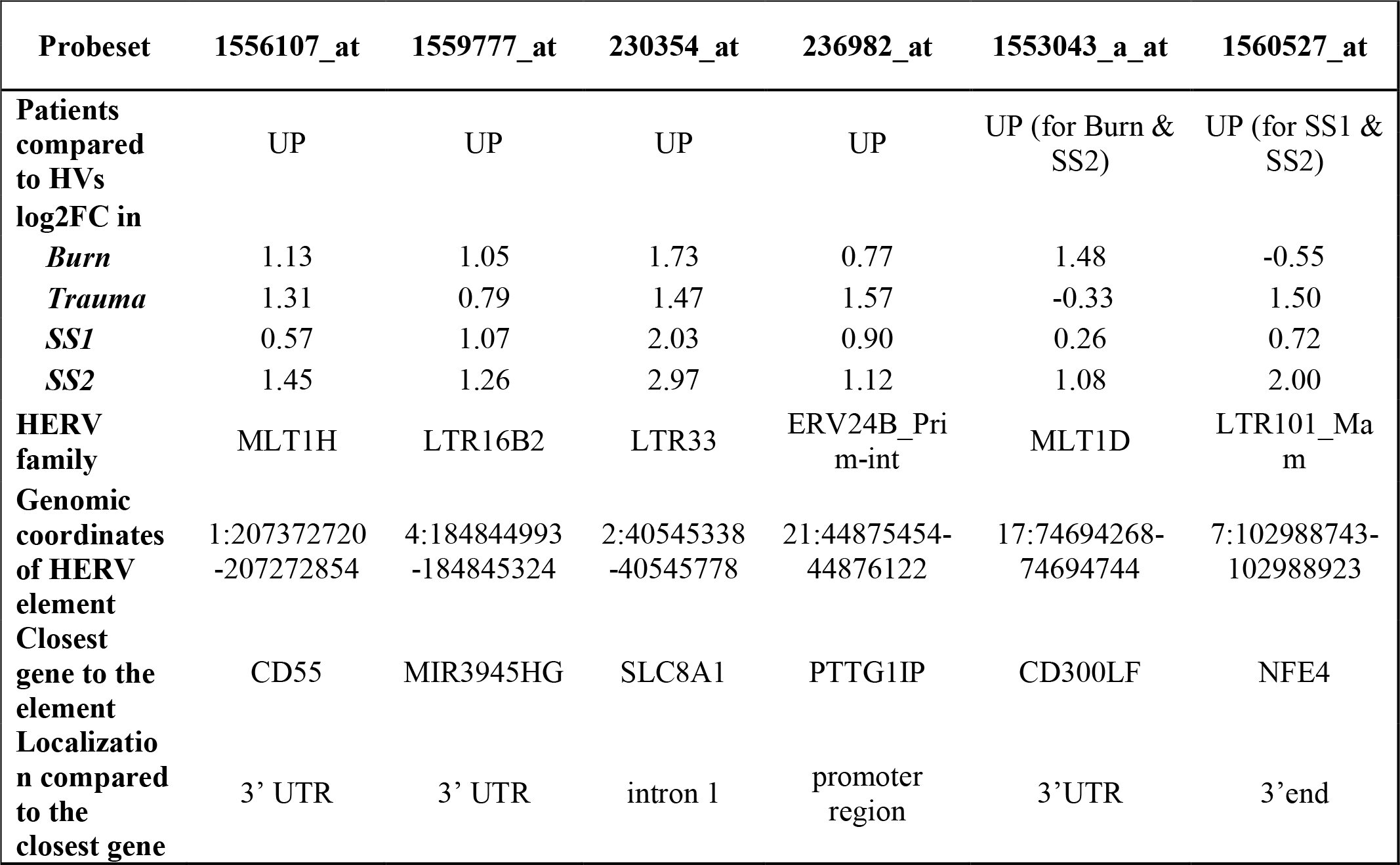
Characteristics of the 6 probesets of interest.

Moreover, we selected 2 other probesets of interest, 1553043_a_at and 1560527_at (Figure 6C). The first one targets a MLT1D HERV located in the 3’UTR of *CD300LF*. It was up-modulated in burn and SS2 cohorts. It had a strong up-modulation at D1 in burn patients compared to HV, decreasing over the first week towards HV expression level at D7. The second one targets a LTR101_Mam HERV located in a 3’UTR of a processed transcripts of *NFE4* gene. It was differentially expressed in the 2 septic shock cohorts. This probeset had the highest log2FC among the 5 septic shock-specific modulated probesets.

To validate these transcriptional HERV modulations, we designed primers on the 6 described HERV loci above and on nearby genes by RT-qPCR (Table S2). For each targeted region, we made multiple RT-qPCR designs. We identified several distinct patterns of expression comparing HERVs and nearby genes: (i) for *PTTG1IP* and *MIR3945HG* regions, we observed no or low signal from the HERV loci (data not shown), (ii) for SLC8A1 (Figure 7) and *NFE4* (Figure 8) regions, we observed a high signal from HERVs elements, but no or lower signal on the genes, (iii) for *CD55* (Figure 9) and *CD300LF* (Figure 10) regions, we observed a middle or high signal from both HERV loci and genes.

**Figure 7:**
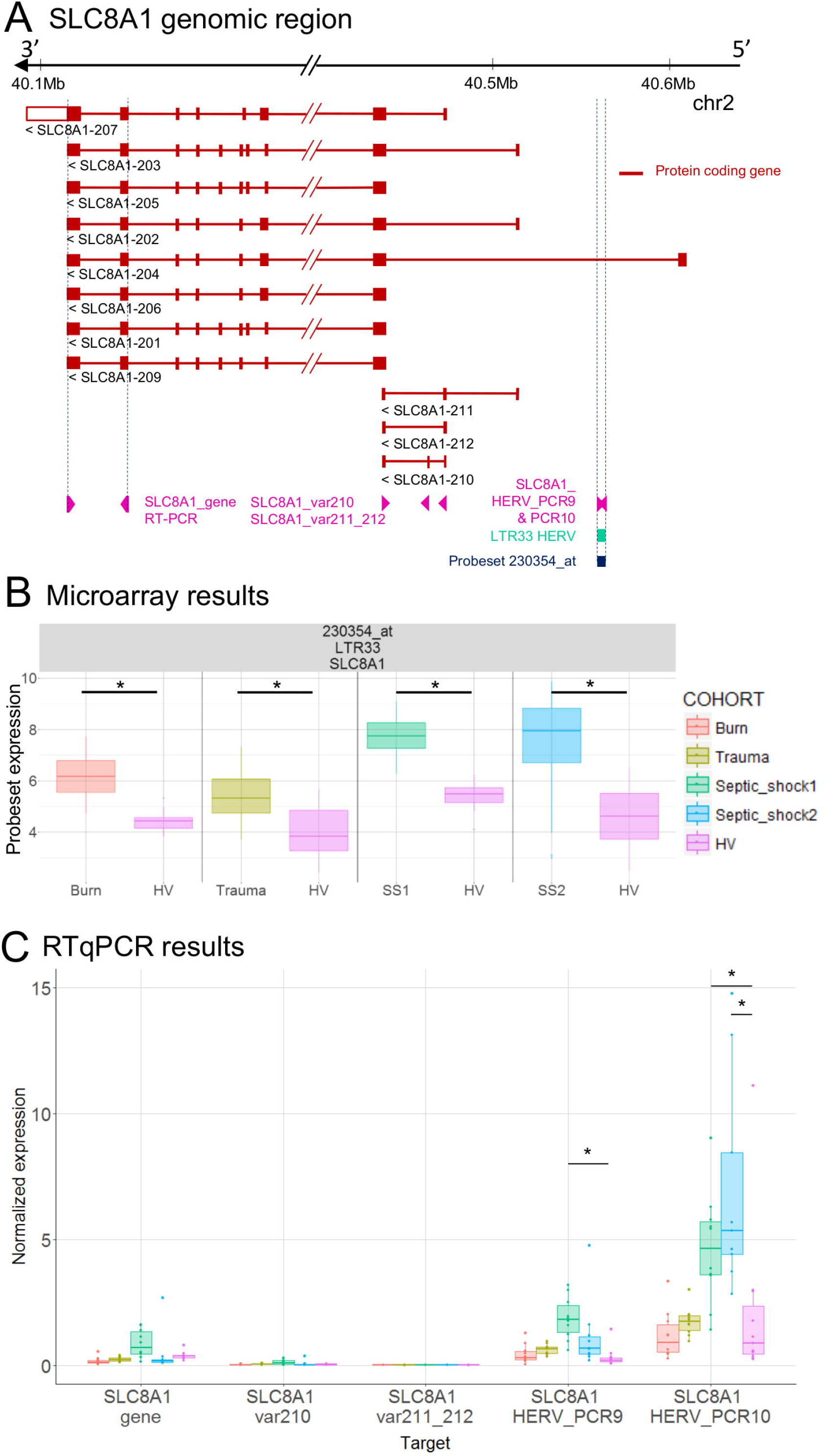
LTR33 HERV and *SLC8A1* gene expression. (A) *SLC8A1* genomic region, with the position of HERV in green, probeset in dark blue and PCR designs in purple. **(B)** Expression levels of the HERV loci from microarray, in HV and patients at D1. **(C)** Expression levels of specific transcripts by RT-qPCR, as described in A, in HV and patients at D1. Expression levels (copy number / μl) were normalized with reference gene (*HPRT1*). Boxes are color-coded by cohort. Statistically significant difference with HV is marked by * (Wilcoxon signed rank test, p-value <0.05).

**Figure 8:**
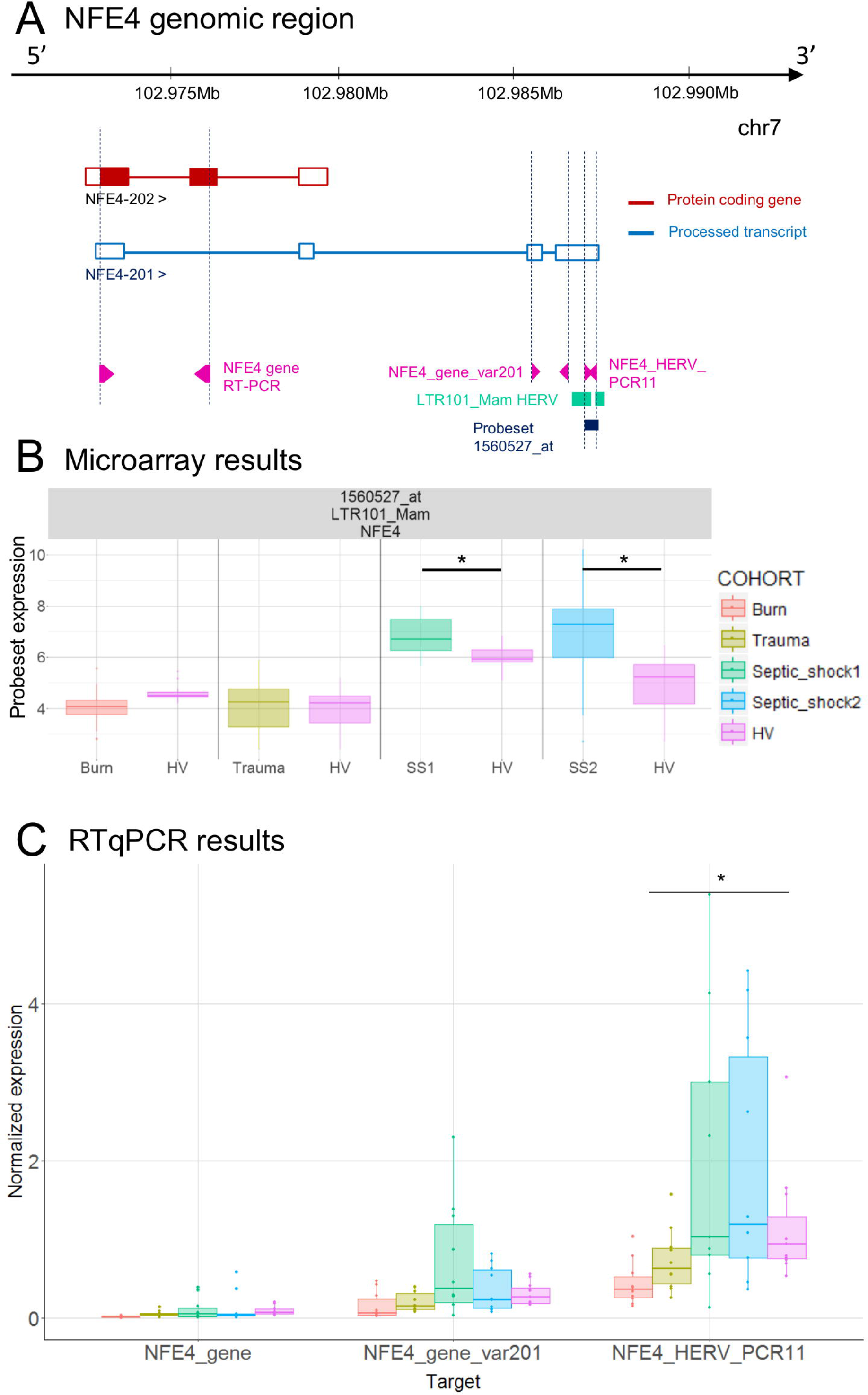
LTR101_Mam HERV and *NFE4* gene expression. **(A)** *NFE4* genomic region, with the position of HERV in green, of probeset in dark blue, of PCR designs in purple. **(B)** Expression levels of the HERV loci from microarray, in HV and patients at D1. **(C)** Expression levels of specific transcripts by RT-qPCR, as described in A, in HV and patients at D1. Expression levels (copy number / μl) were normalized with reference gene (*HPRT1*). Boxes are color-coded by cohort. Statistically significant difference with HV is marked by * (Wilcoxon signed rank test, p-value <0.05).

**Figure 9:**
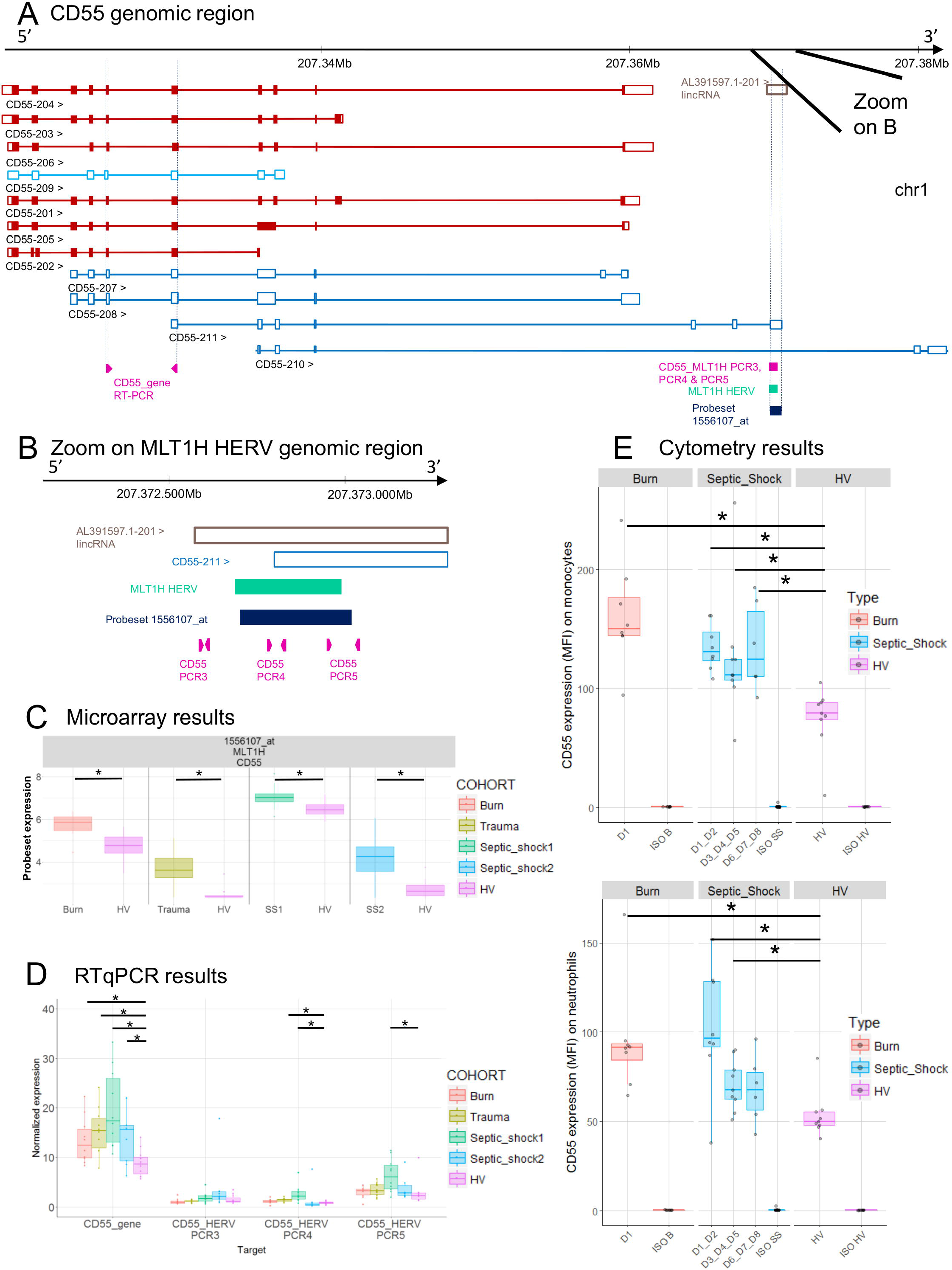
*CD55* associated HERV. **(A)** *CD55* genomic region, with the positions of HERV in green, of probeset in dark blue, of PCR designs in purple. **(B)** Zoom in genomic region of HERV showing PCR designs in detail. **(C)** Expression levels of the HERV loci from microarray, in HV and patients at D1. **(D)** Expression levels of specific transcripts by RT-qPCR, as described in A and B, in HV and patients at D1. Expression levels (copy number / μl) were normalized with reference gene (*HPRT1*). Boxes are color-coded by cohort. **(E)** Protein expression levels (MFI), on monocytes (left) and neutrophils (right) from 8 burn patients (red), 11 septic shock patients (blue) and 9 HV (purple). Columns ISO B, ISO SS and ISO HV correspond to isotypes for burn, septic shock and HV respectively. Statistically significant difference with HV is marked by * (Wilcoxon signed rank test, p-value <0.05).

**Figure 10:**
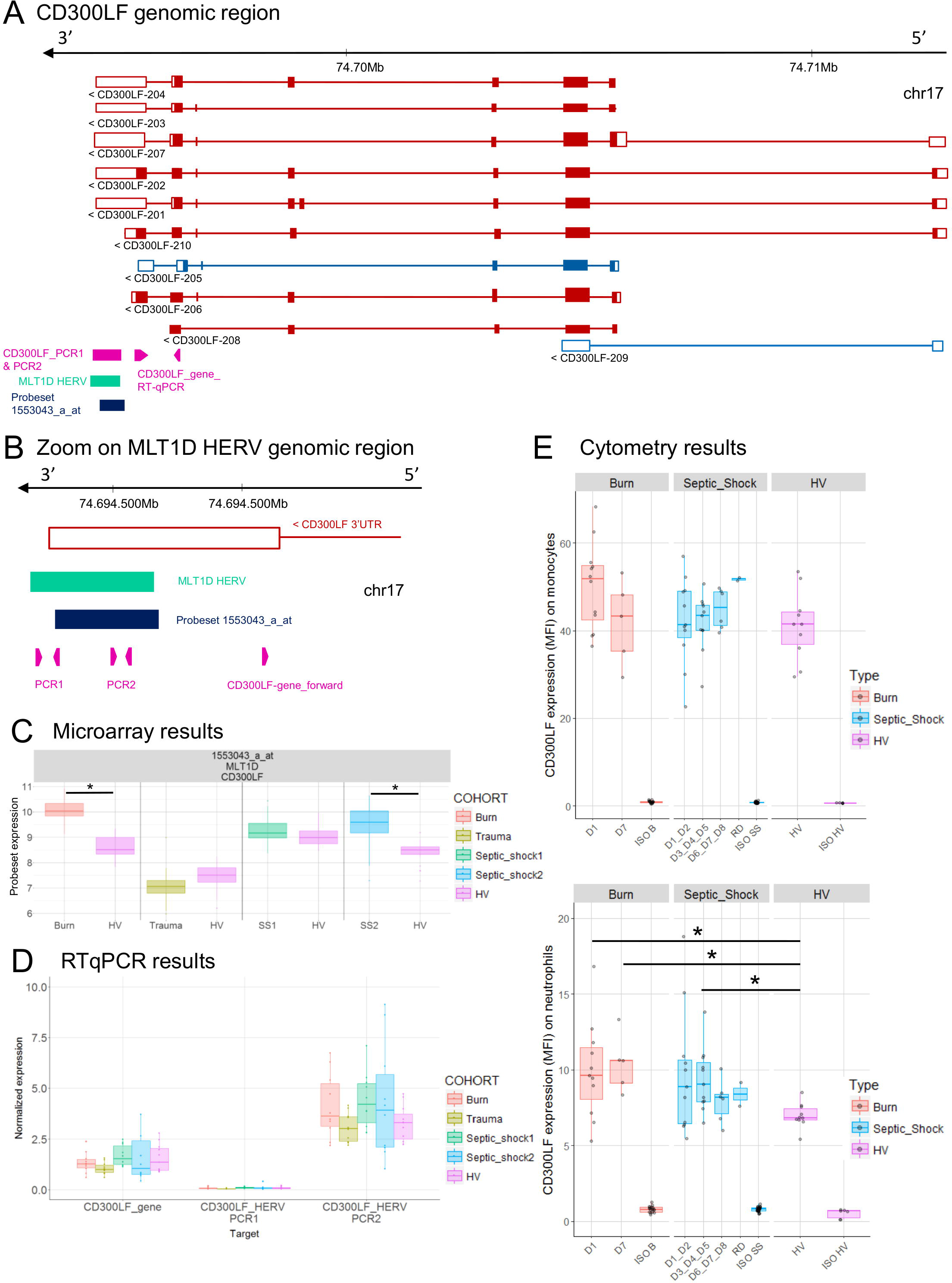
*CD300LF* associated HERV. **(A)** *CD300LF* genomic region, with the positions of HERV in green, of probeset in dark blue, of PCR designs in purple. **(B)** Zoom in genomic region of HERV showing PCR designs in detail. **(C)** Expression levels of the HERV loci from microarray, in HV and patients at D1. **(D)** Expression levels of specific transcripts by RT-qPCR, as described in A and B, in HV and patients at D1. Expression levels (copy number / μl) were normalized with reference gene (*HPRT1*). Boxes are color-coded by cohort. **(E)** Protein expression levels (MFI), on monocytes (left) and neutrophils (right) from 14 burn patients (red), 11 septic shock patients (blue) and 10 HV (purple). Columns ISO B, ISO SS and ISO HV correspond to isotypes for burn, septic shock and HV respectively. Statistically significant difference with HV is marked by * (Wilcoxon signed rank test, p-value <0.05)

To better interpret the results, we extracted from Ensembl the genome annotation and showed in genomic context, the microarray and the RT-qPCR results of *SLC8A1* (Figure 7), *NFE4* (Figure 8), *CD55* (Figure 9) and *CD300LF* (Figure 10) regions. *SLC8A1* has 11 known transcripts. All but one are located in 3’ of the LTR33 HERV element targeted by the 230354_at probeset, which is located in the first intron of SLC8A1-204 transcript (Figure 7A). The up-modulation of the LTR33 element in septic shock patients observed on microarray was confirmed by RT-qPCR (Figure 7B, Figure 7C). The up-modulation observed for other cohorts was not confirmed by RT-qPCR. The gene *SLC8A1* was not expressed in patients or HV, as seen on various microarray probesets and confirmed by RT-qPCR (SLC8A1_gene, var210, var211_212).

*NFE4* gene has 2 transcripts (Figure 8A) and only one is coding for a protein (NFE4-202). The LTR101_Mam HERV element, targeted by the 2560527_at probeset, is located in 3’UTR of NFE4-201, the non-protein-coding transcript. Although the same trends are observed between microarray and RT-qPCR, the up-modulation of the LTR101_Mam element observed in septic shock patients with microarray was not statistically significant in RT-qPCR, (Figure 8B, Figure 8C). There was low or no signal on designs targeting gene transcripts (NFE4_gene and NFE4_gene_var201).

*CD55* gene has 11 transcripts. The MLT1H HERV element, targeted by the 1556107_at probeset, is located in the 3’UTR of CD55-211 transcript (Figure 9A). The HERV element overlaps the 3’UTR of transcript CD55-211 and a long intergenic noncoding RNA (lincRNA, a class of long transcribed RNA molecules longer than 200 nucleotides and not coding for proteins) (Figure 9B). The up-modulation of MLT1H saw with microarray in the 4 cohorts was partially confirmed with RT-qPCR on trauma and septic shock cohorts (Figure 9C, Figure 9D). The designs targeting MLT1H or close neighborhood (PCR3, 4 and 5) presented the same profile, with a significant difference in septic shock and trauma cohorts compared to HV (PCR4). The design targeting the gene showed also up-modulation of *CD55* and a very high absolute normalized expression in patients compared to HV (Figure 9D). (Of note 1555950_a_at probeset, targeting most of *CD55* transcripts, was also up-modulated in patients, and with a high expression level (data not shown)). We also confirmed by flow cytometry on monocytes and neutrophils that *CD55* expression was higher in patients than in HV, confirming an up-modulation at the protein level in patients(Figure 9E).

The MLT1D HERV element, targeted by the 1553043_a_at probeset is located in 3’UTR of CD300LF-201, 202, 203, 204 and 207 protein-coding transcripts (Figure 10A). We made several RT-qPCR designs, targeting either the HERV locus only (PCR1) or both HERV and 3’UTR of *CD300LF* (PCR2, Figure 10B). The up-modulation seen in burns and septic shock 2 cohorts on microarray was not confirmed by RT-qPCR, neither for gene nor for HERV designs (Figure 10D). PCR1 showed no signal at all. PCR2 design showed a slight higher expression level in burn and septic shock cohorts compared to HV. We also confirmed an higher expression at the protein level by flow cytometry on neutrophils in burn and septic shock patients, compared to HV (Figure 10E). In monocytes, protein level in burn at D1 seemed slightly higher than HV.

## Discussion

We took advantage of previous microarray analyses on four cohorts of severely injured patients to assess the modulation of HERV transcriptome in acute inflammation. We showed that several loci were expressed and modulated after acute injury. Surprisingly, a large majority among the modulated HERVs were down-modulated in patients compared to HV, whereas a global and massive gene up-modulation has been observed after severe injuries (Xiao et al. 2011).

Five HERVs were modulated in patients compared to HV in all four datasets and 16 HERVs in at least 3 datasets, suggesting a similar inflammatory triggered modulation in all models. We validated expression profiles by RT-qPCR on 6 regions, allowing us to explore more precisely the modulation pattern of the HERVs and the neighbor genes. Interestingly, all these 6 HERVs have detected signals in RNAseq experiments from lymphoid cells and whole blood datasets (Ensembl Rnaseq tracks, (Aken et al. 2017)). Some authors already focused on HERV detection in blood of burn patients using pan-family RT-PCRs (Y.-J. Lee et al. 2013; K.-H. Lee et al. 2014). However, very few data are available in human diseases for specific loci. No study had yet evaluated the expression of HERVs in acute inflammatory contexts by using multiple cohorts with different types of inflammatory injuries.

Several groups showed that huge epigenetic modifications occur after acute inflammation, regulating transcriptional profiles in the immune system, especially in sepsis (J. L. G. Gimenez et al. 2016; Saeed et al. 2014). These epigenetic modifications may explain the polarization profiles such as tolerance or trained immunity, observed after various stimulations of innate cells (Saeed et al. 2014). We hypothesized and confirmed *in vivo* that other elements than genes, especially HERVs which are known to be tightly controlled by epigenetic modifications (Daskalakis et al. 2018), might be modulated in acute inflammatory situations. This has also been demonstrated in other pathophysiological contexts such as cancer (J. Gimenez et al. 2010; Pérot et al. 2015; Lamprecht et al. 2010; Beyer et al. 2016), where global epigenetic modifications are also observed (Chiappinelli et al. 2015; Groh and Schotta 2017).

Interestingly in cancer, epigenetic modifications that gave access to HERV cis sequences through open chromatin, have also revealed a very role in pathophysiology (Lamprecht et al. 2010; Cohen, Lock, and Mager 2009; Mager et al. 1999). Indeed, by providing alternative promoter sequences to classical protein coding genes, these epigenetic modifications explain part of the ectopic expression of myeloid-growth factor receptors in lymphoid cells (Lamprecht et al. 2010). This underlines how HERV elements, in particular their LTRs, could modulate gene expression and the host immune response to injury. In our study, the four commonly modulated HERVs were LTRs located nearby genes related to the immune response. In several cases (*NFE4*, *CD300LF*), we found a polyadenylation signal (AAUAAA) provided by the HERV LTR in 3’ of some of the alternative transcripts of the genes. The case of *CD300LF* is interesting as this protein acts as an inhibitory receptor for myeloid cells (Alvarez-Errico et al. 2004). The LTR might stabilize specific transcripts and enhance expression of CD300LF protein, which we confirmed by flow cytometry in severe burn patients early after admission. This up regulation might participate in the compensatory anti-inflammatory response. The precise understanding of the mechanisms through which specific HERV LTRs might impact immune gene expression is not possible in such translational research setting with patient samples. This will require in the future *in vitro* experimental models to validate and understand our observations.

Our RT-qPCR validation assays also showed inter-individual variability and underlined that exploring such repertoire of our genome, repetitive sequences, may face specificity issues, and will require specific tools. Indeed, as a first attempt, we used commercial microarray where probesets were not initially designed to target HERV elements. Moreover, as the probesets targeting HERVs were initially supposed to target conventional genes, the majority of explored HERVs are close to or within a gene. To better understand HERV expression in these settings, targeting HERVs localized far from genes seems important. Until now, the lack of tool made difficult the exploration of HERV expression. It would be interesting to reproduce these analyses, with a more exhaustive technology designed to specifically target HERVs, like the HERV-V3 Affymetrix microarray we recently published (Becker et al. 2017), or even RNAseq. It will allow us to better describe the whole HERV transcriptome modulation and understand the putative global role of HERV in the host response.

Finally, it would be of importance to take into account HERV expression in further blood transcriptome analyses, especially in such acute inflammatory contexts, to better understand HERV expression during host response. HERVs could be good markers of the different phases after important inflammatory shocks and could even become potential therapeutic targets if their functional role on host-response is confirmed.

To conclude, we showed for the first time that specific HERV loci are transcribed in whole blood of ICU patients. Our design allowed us to identify specific transcriptional signatures of HERVs elements, *in vivo*, linked to the acute inflammatory response. Moreover, the similarities observed in three models of acute injuries suggest common regulatory mechanisms and a specificity of the observed modulation. We also unravel the potential regulatory role of these elements within the host immune response. Further studies are needed to better understand such mechanisms and how HERVs may contribute to the pathophysiology of the host immune response, a key part of the pathophysiology of sepsis.

### List of abbreviations

HERV: Human endogenous retrovirus; LTR: Long Terminal Repeats; PAMP: pathogen-associated molecular pattern; LPS: Lipopolysaccharide; PMA: phorbol-12-myristate-13-acetate; HV: healthy volunteers; ICU: Intensive Care Unit; TBSA: Total Burn Surface Area; ABSI: Abbreviated Burn Severity Index; ISS: Injury Severity Score; SAPSII: Simplified Acute Physiology Score II; MFI: Medians of Fluorescence Intensity;

## Acknowledgements

The authors would like to gratefully thank Maria-Paola Pisano, Marie-Angélique Cazalis, Boris Meunier, Julie Mouillaux and Estelle Peronnet for their kind advices. They also thank all clinical research assistant for the collection of blood samples, especially Hélène Vallin and Valérie Cerro. Finally they gratefully thank Anne Portier and Marie-Angélique Cazalis for all experiments made on samples.

## Author Contributions Statement

OT and JT designed the project, performed the analyses and wrote the paper. CJ and FV performed cytometry experiments. EC performed RT-qPCR validations. ML, AC, BA, TR recruited patients in the various cohorts. OT, MM, FV, FM, JT read and discussed the manuscript. All authors drafted or revised critically the manuscript for important intellectual contents. All authors read and approved the final manuscript.

## Data availability statement

Microarray expression data has been deposited on NCBI Gene Expression Omnibus and are accessible through GEO accession numbers GEO:GSE77791, GEO:GSE57065 and GEO:GSE95233. Data from microarray experiment for trauma cohort are available at Hospices Civils de Lyon – bioMérieux – UCBL1 “Pathophysiology of Injury Induced Immunosuppression”, Groupement Hospitalier Edouard Herriot, France.

